# Dopaminergic and opioidergic regulation of implicit hedonic facial reactions during anticipation and consumption of social and nonsocial rewards

**DOI:** 10.1101/832196

**Authors:** Sebastian Korb, Sebastian J. Götzendorfer, Claudia Massaccesi, Patrick Sezen, Irene Graf, Matthäus Willeit, Christoph Eisenegger, Giorgia Silani

**Author notes:** Correspondence: Sebastian Korb, Department of Psychology, University of Essex, Colchester, CO4 3SQ, UK., Giorgia Silani, Department of Clinical and Health Psychology, University of Vienna, Liebiggasse 5, 1010 Vienna, Austria.

## Abstract

The observation of animal hedonic orofacial and behavioral reactions has played a fundamental role for the identification of a dopaminergic motivational, and an opioidergic hedonic component of reward. Translation to humans remains difficult, however, as human research has struggled to adopt a similar operationalization of reward. Here, we investigated the neurochemical basis of hedonic facial and behavioral reactions to different types of rewards in healthy adult volunteers, by pharmacologically reducing dopaminergic and opiodergic receptor-specific action. Subjective ratings, physical effort, and facial reactions to matched primary social (affective touch) and nonsocial (food) rewards were assessed. Both drugs resulted in reduced physical effort and increased negative facial reactions during reward *anticipation*, but only opioidergic manipulation caused reduced positive facial reactions during reward *consumption*. This suggest that facial reactions during anticipated and experienced pleasure rely on partly different neurochemical systems, providing novel evidence in support of existing theoretical models of reward.

## Introduction

Rewards are salient stimuli, objects, events, and situations that induce approach and consummatory behavior by their intrinsic relevance for survival or because experience has taught us that they are pleasurable ^1^. Rewards can be parsed, at the psychological, neurophysiological, and neurochemical level, into the main components *wanting* (the motivation to mobilize effort to obtain a reward), *liking* (the hedonic response evoked by its consumption) ^2–4^. This conceptual division is paralleled in cognitive theories of economic decision making ^5,6^ that similarly distinguish between *decision utility* (how much the value attached to an outcome determines its choice or pursuit), and *experienced utility* (referring to the subjective hedonic experience generated by an outcome). Today, our understanding of wanting and liking rests on 30 years of animal research, and on preliminary confirmatory findings in humans, and the parsing of rewards into its subcomponents has been shown to have important implications for affective and addictive disorders, including substance abuse and schizophrenia ^7,8^.

Wanting is mainly linked to the mesolimbic dopaminergic system, and is dissociable from liking, which instead relies on the opioidergic system, as suggested for example by the “taste reactivity test”, a method to assess eating-related pleasure by observing facial and bodily reactions of animals and human infants to palatable and aversive tastes ^9,10^. For instance, neither pharmacological disruption, nor extensive lesion of dopaminergic neurons affect facial liking reactions (e.g. relaxed facial muscles and licking of the lips) to consumption of sweet foods in rats ^11,12^, but increased mesolimbic dopamine release induced by electric stimulation of the hypothalamus results in greater wanting to eat (e.g. food intake) without modulating liking ^13^. On the other hand, hedonic reactions to sensory pleasure are amplified by opioidergic stimulation of various “hedonic hotspots” of the brain, including the nucleus accumbens (NAc) shell and parts of the limbic system ^3^. In addition, the opioidergic system partly also affects wanting by modulating the effects of dopamine in the NAc ^14^. Indeed, injections of µ and δ opioid receptor agonists have been shown to increase food approach and feeding behavior, especially for palatable and high-energy foods ^15^, which suggests that opioids primarily affect wanting through liking.

Evidence of similar neurochemical parsing of reward processing in humans is mainly derived from research in clinical populations, and a handful of recent studies in healthy volunteers. For example, stimulation of D2/D3 receptors through dopamine agonists can induce compulsive medicament use, gambling, shopping, hypersexuality, and other addictive activities in some patients with Parkinson’s disease ^16,17^. These behaviors, which correspond to strong urge-like wanting without changes in subjective liking, are accompanied by altered activations in the ventral striatum and prefrontal cortex, which however normalize when patients are off dopaminergic medication. In healthy volunteers, dopamine D2/D3 receptor blockade with amisulpride disrupts the motivation to gain immediate rewards in both a pavlovian-instrumental-transfer task and a delay discounting task ^18^, and reduces the rewarding value of prosocial decisions in women ^19^. Administration of µ opioid receptor agonists increases the subjective pleasantness of the most palatable food option available ^20^, and both subjective feelings of wanting and liking of the most attractive opposite sex faces ^21^. In contrast, the non-selective opioid receptor antagonist naloxone decreases subjective pleasure associated with viewing erotic pictures and reduces the activation of reward related brain regions such as the ventral striatum ^22^.

In spite of the progress made, human and animal research on reward processing remain difficult to compare, as human research has struggled to adopt an operationalization that resembles the one used in animal research, e.g. measuring behavior and facial reactions instead of relying on subjective verbal report ^8,23^. Indeed, while the decision utility (wanting) of a reward is easily inferred from observed choices, such as purchasing a good, or the effort mobilized to obtain it, the concomitant experienced utility (liking) is more challenging to measure objectively. In human newborns, juvenile monkeys, and adult rats the consumption of food rewards elicits powerful and distinctive facial reactions ^9,24,25^. The taste reactivity test ^9^ has indeed become the gold standard to assess hedonic consummatory pleasure in animal models. Furthermore, facial reactions to rewards are not restricted to the consummatory phase, but can also be observed during the anticipation of a reward. Indeed, following Pavlovian conditioning, animals show hedonic facial reactions to cues, which they learned to associate with the delivery of an unconditioned taste stimulus ^26^. In contrast to the facial reactions occurring during reward consumption, facial reactions to the conditioned stimuli reflect the prediction of pleasure from a future reward (i.e. anticipated pleasure) ^23^. Whether hedonic facial reactions to anticipated or consumed rewards are neurochemically regulated in a similar way has, to our knowledge, never been investigated.

Hedonic facial reactions to pleasant tastes and other types of reward are more subtle in human adults, and have only started to be investigated using facial electromyography (fEMG) ^27–31^. Recently, we have shown ^27^ that the anticipation and consumption of preferred food rewards results in the relaxation of the main frowning muscle corrugator supercilii (CS), and that the experience of pleasant social touch elicits activation of the main smiling muscle, the zygomaticus major (ZM). The latter result only emerged from explorative analyses, but others have also reported ZM contraction and CS relaxation in response to pleasant social touch ^28–30^. Extant research thus suggests that human adults relax the CS and activate the ZM during both the anticipation and the consumption of different types of pleasurable stimuli, although differences between types of rewards may also exist. How these hedonic facial reactions rely on the dopaminergic and opioidergic systems, and whether they are differently modulated depending on the type of reward announced and delivered, is currently unknown. Importantly, establishing the neurochemical basis of different aspects of reward in humans, by using translational tasks allowing better cross-species comparison, is expected to contribute to our understanding – and ultimately treatment – of reward anomalies occurring in several neuropsychiatric disorders like schizophrenia, depression, and autism ^8^.

To fill this gap of knowledge, we pharmacologically manipulated the dopaminergic and opioidergic systems in adult humans and measured both explicit (subjective ratings, physical effort) and implicit (fEMG) reactions during anticipation and consumption of social and nonsocial rewards. Sweet milk with different concentrations of chocolate flavor served as nonsocial rewards. Gentle caresses to the forearm (typically referred to as “social touch” or “affective touch” in the literature), delivered by a same-sex experimenter at different speeds and resulting in different levels of pleasantness ^32–34^, served as non-sexual social rewards. Importantly, these can both be considered to be primary rewards (i.e. a biological preparedness can be expected), and we conducted extensive prior work to select stimuli of similar magnitude across reward type ^27^. In addition, trial-by-trial ratings during this experiment confirmed that the social and nonsocial rewards used had comparable reward magnitudes for our participants. Similarly to effort-related choice tasks used in animals ^35^ and humans ^36,37^, participants had the choice between exerting physical effort to obtain a greater reward, and exerting less or no physical effort to obtain a smaller reward. In each trial of the experiment (Fig. 1), rewards with high or low value were announced and participants were asked to rate how much they wanted the reward, and to exert physical effort (squeezing of an individually thresholded hand-dynamometer) that was directly converted in the probability to receive the announced reward, or, alternatively, the least liked reward. The reward obtained (either the announced ‘high’ or ‘low’ reward, or the ‘verylow’ reward following insufficient effort), was subsequently delivered and its liking was measured with subjective ratings.

**Fig. 1:**
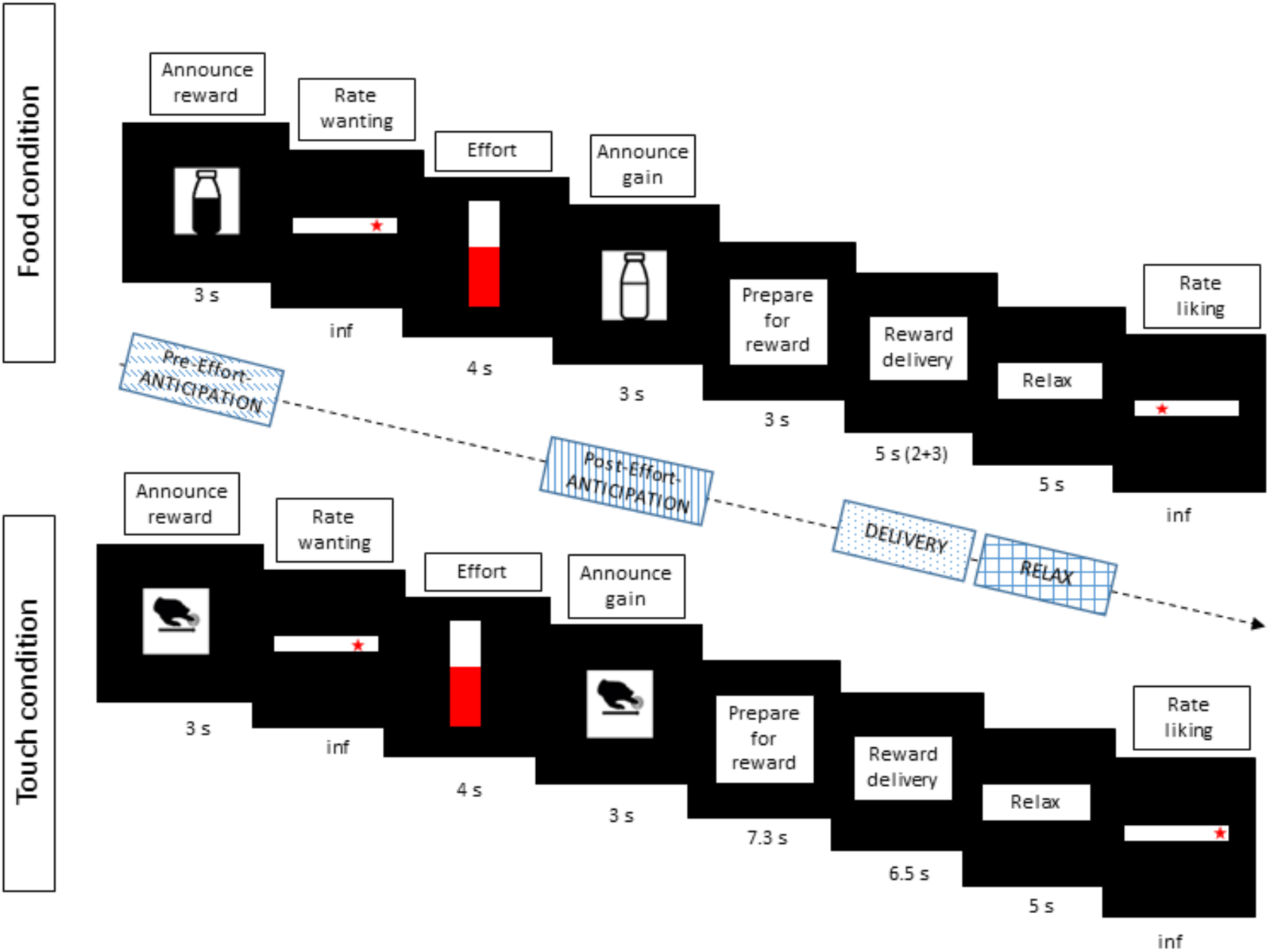
Main elements in each trial for the Food and Touch conditions. Before the main task, participants individually ranked three reward levels per condition by means of liking-ratings. In the main task (here depicted), one of the two most liked rewards (high and low) was announced at the beginning of each trial. The probability of obtaining the announced reward was determined linearly by participants’ hand-squeezing effort, and indicated in real-time. The gained reward (which was either the one announced at the beginning of the trial, or the least-liked verylow reward if squeezing was not sufficient) was then announced and delivered. To assess reward anticipation, EMG data was analyzed during the Pre-Effort anticipation period (3 sec) at the beginning of the trial, when a possible reward was announced, as well as during the Post-Effort anticipation period (3 sec announcement) preceding reward delivery. To investigate reward consumption, EMG data was analyzed during reward Delivery (5 sec in the Food, and 6.5 sec in the Touch condition), and in the immediately following Relax phase (5 sec). “Inf” (under ratings) symbolizes that ratings slides stayed on screen indefinitely, or until participants’ button press. For a complete representation of all trial elements see Fig. S1 in the Supporting Information.

Implicit hedonic facial reactions during the anticipation of the reward as well as during and immediately after its delivery were recorded together with subjective ratings.

The role of the dopaminergic and opioidergic systems was investigated using oral administration three hours before the task of the highly selective D2/D3 dopamine receptor antagonist amisulpride (400 mg), the non-selective opioid receptor antagonist naltrexone (50 mg), or placebo, in a randomized, double-blind, between-subject design in 131 healthy volunteers (group sizes were 42, 44, and 45 respectively for amisulpride, naltrexone, and placebo). Sample sizes were chosen based on previous work employing the same compounds and doses ^18^.

Adopting a translational approach and operationalizing reward processing in humans in a way that makes it comparable to animal research (e.g. measuring real effort and hedonic facial responses), we investigated two fundamental yet unresolved research questions: 1) to what extent do motivational and hedonic implicit and explicit responses to rewards rely on the dopaminergic and opioidergic systems in humans, and 2) do social (touch) and nonsocial (food) rewards share the same neurochemical basis in humans.

We made the following hypotheses based on the literature. First, because liking relies heavily on the opioidergic but not the dopaminergic system, subjective ratings of liking, and hedonic facial reactions during and after reward consumption, were expected to be reduced after administration of the opioid antagonist naltrexone, compared to placebo, but not after administration of the dopamine antagonist amisulpride. Second, because wanting is believed to be regulated by the dopaminergic and indirectly also by the opioidergic systems, we expected subjective ratings of wanting, and physical effort applied to obtain the announced reward, to be reduced after administration of both the D2/D3 receptor antagonist amisulpride, and naltrexone. Third, because facial responses during reward anticipation – previously shown to occur to learned cues for rewards in rats^26^ and humans^27^ – may reflect anticipated pleasure during a period commonly associated with wanting, they were expected to be affected by naltrexone, as well as by amisulpride, compared to placebo. Finally, based on on fEMG results showing similar facial reactions to different types of rewards ^27–30^, and on evidence from neuroimaging studies that supports the ‘common currency hypothesis’ of reward processing ^3,38^, we expected the same pattern of results for food and touch rewards.

## Results

### Matching of drug groups

Given that the type of reward received in each trial depended on both the announcement cue at the beginning (high or low) and the force exerted to obtain it (verylow rewards were only obtained following insufficient effort), we tested for differences across groups. The number of trials with high, low, or verylow rewards obtained did not differ across groups, as shown by a linear mixed effects model (LMM) with number of trials as dependent variable, the fixed effects Condition (social, nonsocial), Drug (amisulpride, naltrexone, placebo), and Reward Type (high, low, verylow), and by-subject random intercepts. Only a significant main effect of Reward Type was found (*F*(2, 763) = 27.84, *p* < .001), due to a greater number of high (*M* = 16.53, *SD* = 2.87) than low (*M* = 14.93, *SD* = 3.46) and verylow (*M* = 8.67, *SD* = 4.93) trials received, across all three drug groups.

Moreover, groups of participants did not differ (Table 1) in their average age, BMI, maximum voluntary contraction (MVC) of the hand dynamometer measured right before the main task, nor did they differ in their mood, which was measured with the Positive and Negative Affect Schedule (PANAS) ^39^ at time of pill intake (T1) and three hours later (T2) allowing us to exclude differences across groups as possible confounds of our results.

**Table 1:** participant characteristics. BMI = Body Mass Index; MVC = Maximum Voluntary Contraction; PANAS = Positive and Negative Affective Schedule; M = Mean

|  | **Amisulpride** | **Naltrexone** | **Placebo** | Group differences |
| --- | --- | --- | --- | --- |
| N (Male, Female) | 42 (14, 28) | 44 (14, 30) | 45 (15, 30) |  |
| Age M (SD) | 23.7 (4.1) | 22.9 (2.8) | 23.1 (3.7) | $t = -0.73, p = 0.46$ |
| BMI M (SD) | 22.7 (2.5) | 23.0 (2.3) | 22.2 (2.5) | $t = -0.99, p = 0.32$ |
| MVC M (SD) | 211.9 (86.3) | 208.7 (81.8) | 215.3 (73.1) | $t = 0.19, p = 0.85$ |
| PANAS pos T1 M (SD) | 30.5 (5.4) | 29.7 (7.3) | 29.4 (6.7) | $t = -0.8, p = 0.42$ |
| PANAS neg T1 M (SD) | 12.1 (3.2) | 14.3 (7.5) | 11.5 (2.1) | $t = -0.7, p = 0.52$ |
| PANAS pos T2 M (SD) | 27.1 (6.3) | 24.7 (8.0) | 26.7 (7.4) | $t = -0.3, p = 0.80$ |
| PANAS neg T2 M (SD) | 10.1 (2.8) | 12.1 (5.5) | 10.5 (0.9) | $t = -0.5, p = 0.58$ |

### Explicit measures: ratings of wanting, ratings of liking and physical effort

Subjective ratings of wanting and liking, as well as effort were each analyzed in separate LMMs with Condition (social, nonsocial), Drug (amisulpride, naltrexone, placebo), and Reward Type (high, low as announced at the beginning of each trial for wanting and effort; high, low, verylow as obtained after the effort phase for liking) as fixed effects, and as random effects intercepts for subjects and by-subject random slopes for the effects of Condition, Reward Type, and their interaction. Significant main and interaction effects are only reported for the factor Drug, as they are most relevant to this study. Please see the Supporting Information for exhaustive documentation of statistical results (see also Fig. S2).

No main or interaction effects with the factor Drug were found for ratings of wanting. Behavioral analyses on effort resulted in a significant Condition X Drug interaction (Fig. 2A, *F*(1, 128.31) = 4.54, *p* = .01) reflecting lower effort in the food condition in the amisulpride (*M* = 74.98, *SD* = 26.57) and naltrexone (*M* = 73.51, *SD* = 24.43) groups compared to the placebo group (*M* = 80.20, *SD* = 22.41), while effort levels were similar across drug groups in the social condition (amisulpride: *M* = 78.34, *SD* = 25.14; naltrexone: *M* = 73.78, *SD* = 23.15; placebo: *M* = 76.11, *SD* = 23.51). A marginally significant Reward Type X Drug interaction (Fig.2B) was also found (*F*(1, 128.50) = 2.97, *p* = .05), which was related to reduced effort for low rewards in the amisulpride (*M* = 71.67, *SD* = 27.60) and naltrexone (*M* = 67.90, *SD* = 24.45) groups compared to the placebo group (*M* = 75.65, *SD* = 23.60), but no such difference can be reported for the high rewards (amisulpride: *M* = 81.60, *SD* = 23.09; naltrexone: *M* = 79.29, *SD* = 21.70; placebo: *M* = 80.63, *SD* = 22.23). All other effects were not significant (all *F* < 0.9, all *p* > .4).

**Fig. 2:**
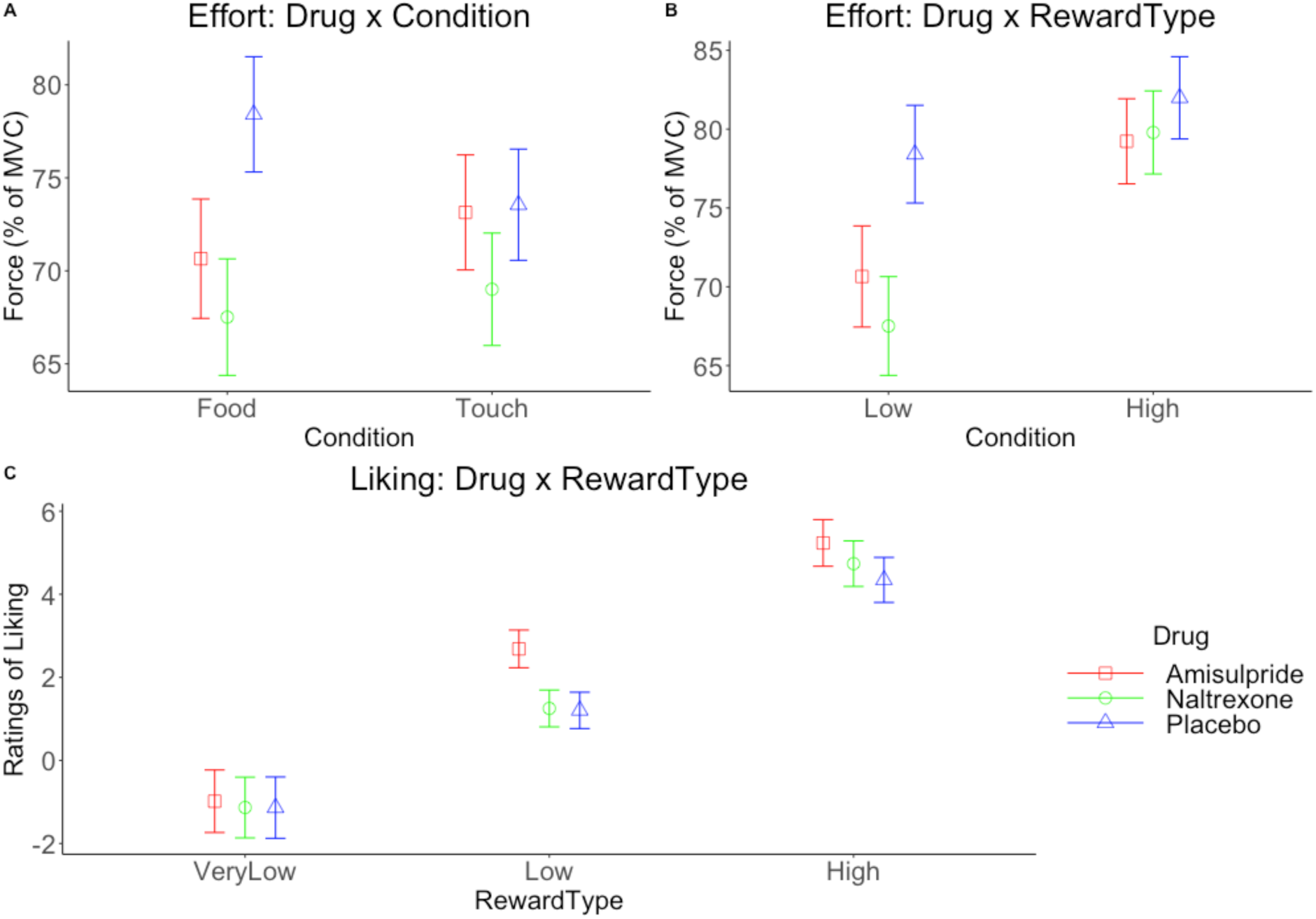
Marginal means (and 95% CIs) for behavioral analyses. Physical effort was reduced in the amisulpride and naltrexone groups compared to placebo in (A) the food condition, and (B) for low rewards. (C) Liking of low rewards was greater in the amisulpride compared to the naltrexone and placebo groups.

The same LMM on the ratings of liking (Fig. 2C) resulted in a significant Drug X Reward Type interaction (*F*(4, 259.62) = 11.07, *p* < .001), reflecting greater liking of low rewards in the amisulpride group (*M* = 2.57, *SD* = 3.78), compared to the naltrexone group (*M* = 1.33, *SD* = 4.35), and the placebo group (*M* = 1.53, *SD* = 4.18).

In summary, the amisulpride and naltrexone groups showed reduced effort to obtain food rewards, and to obtain low rewards of both conditions. The amisulpride group also showed greater liking of low rewards compared to both the naltrexone and placebo groups.

### Implicit measures: facial EMG

Facial EMG analyses were carried out separately for the CS and ZM muscles in four periods of interest (see Fig. 1): “Pre-effort anticipation” during reward announcement at the beginning of each trial, “Post-effort anticipation” during the announcement of the gained reward, “Delivery”, and “Relax”. The EMG of Pre- and Post-Effort anticipation periods was analyzed in relation to ratings of wanting and to effort, as these measures were taken close in time. For the same reason, EMG of the Delivery and Relax periods was analyzed in relation to ratings of liking. To better capture the link between implicit and explicit responses to rewards, and in line with our previous work ^27^, facial EMG of each trial was analyzed in relation to subjective ratings and effort measured during the same trial, as opposed to using a-priori reward levels.

We first investigated if facial EMG reflected subjective ratings of wanting and/or liking and effort, independently of Drug, by regressing the EMG of each muscle (expressed as percentage of the baseline) onto the factors Condition (food, touch) and either Wanting, Liking, or Effort (capitalization to indicate that these are predictors in statistical models). Several main or interaction effects were found, showing that facial EMG was sensitive to changes in reward value, and that it was partly related to participants’ explicit measures of wanting and liking (Figure 3). Activation of the CS muscle was inversely related to Wanting (*F*(1, 138) = 8.00, *p* = .005) and to Effort (*F*(1, 150.13) = 10.33, *p* = .002) in the Pre-Effort anticipation period, and to Liking in the Delivery (*F*(1, 207.82) = 4.65, *p* = .03) and Relax period (*F*(1, 136.85) = 17.03, *p* < .001; but more so in the food condition, as reflected by a Liking X Condition interaction: *F*(1, 140.83) = 6.83, *p* = .009). In contrast, activation of the ZM muscle was positively related to Wanting in the Pre-Effort anticipation period (but only for food, as shown by a Wanting X Condition interaction: *F*(1, 7232.9) = 11.73, *p* < .001), and in the Post-Effort anticipation period (*F*(1, 131.44) = 6.39, *p* = .01). In the Delivery period a trend for a Liking X Condition interaction was observed again mostly for food: *F*(1, 126.70) = 3.2, *p* = .07).

**Fig. 3:**
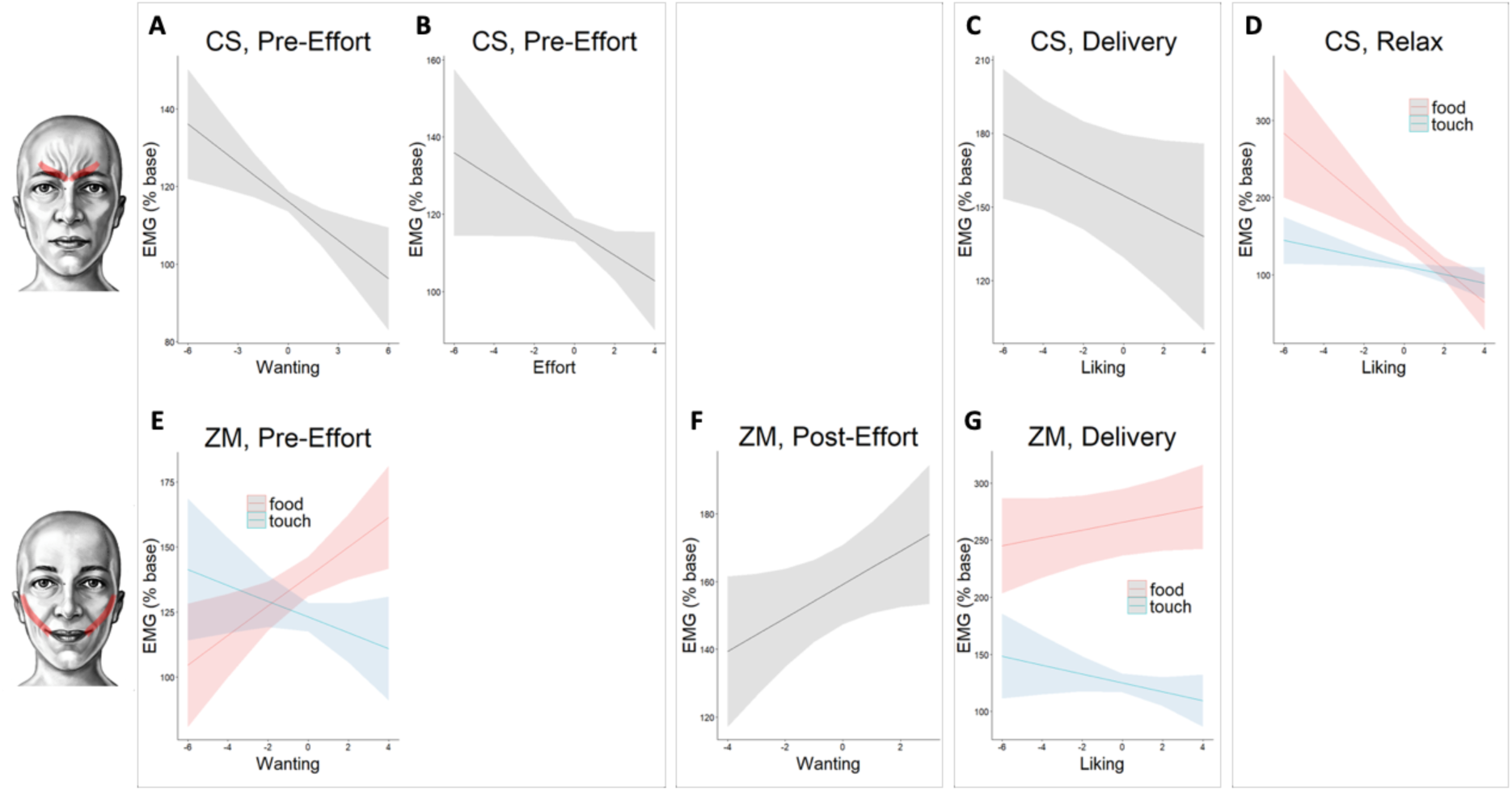
Results (fit lines and and 95% CIs) of analyses investigating the sensitivity of facial EMG to wanting and liking of the administered rewards, independently of drug administration. The CS muscle relaxed with greater (A) wanting and (B) effort in the Pre-Effort anticipation period, and with (C-D) greater liking in the Delivery and Relax periods. The ZM muscle activated with greater wanting in the (E) Pre- and (F) Post-Effort anticipation periods, and (G) in the Delivery period. Some of these effects were restricted to the food condition, as shown by several interactions with Condition (D, E, G).

Next, LMM analyses were carried out to investigate group differences in the facial EMG. These included the factors Condition (food, touch), Drug (amisulpride, naltrexone, placebo), and either Wanting or Effort for the Pre- and Post-Effort anticipation periods, or Liking for the Delivery and Relax periods. Random intercepts for subjects and by-subject random slopes for Condition and Wanting (or Effort/Liking), and their interaction, were included unless indicated otherwise. Only main and interaction effects involving the factor Drug are reported, as they are the most relevant to the study’s hypotheses. Please see the Supporting Information for complete statistical results.

### Pre-Effort anticipation

For the CS muscle by Wanting, a significant Drug X Condition interaction (*F*(2, 128.70) = 4.81, *p* = .009) reflected (Fig. 4) greater CS activation to food than touch in the amisulpride group (*p* = .005; Food: *M* = 119.12, *SD* = 134.43; Touch: *M* = 109.46, *SD* = 76.09) and naltrexone group (*p* < .001; Food: *M* = 120.00, *SD* = 128.19; Touch: *M* = 109.66, *SD* = 89.46), while the placebo group had similar activations across both conditions (*p* = .65; Food: *M* = 110.30, *SD* = 65.72; Touch: *M* = 111.44, *SD* = 78.17). All other effects were not significant (all *F* < 2.2, all *p* > .1).

**Fig. 4:**
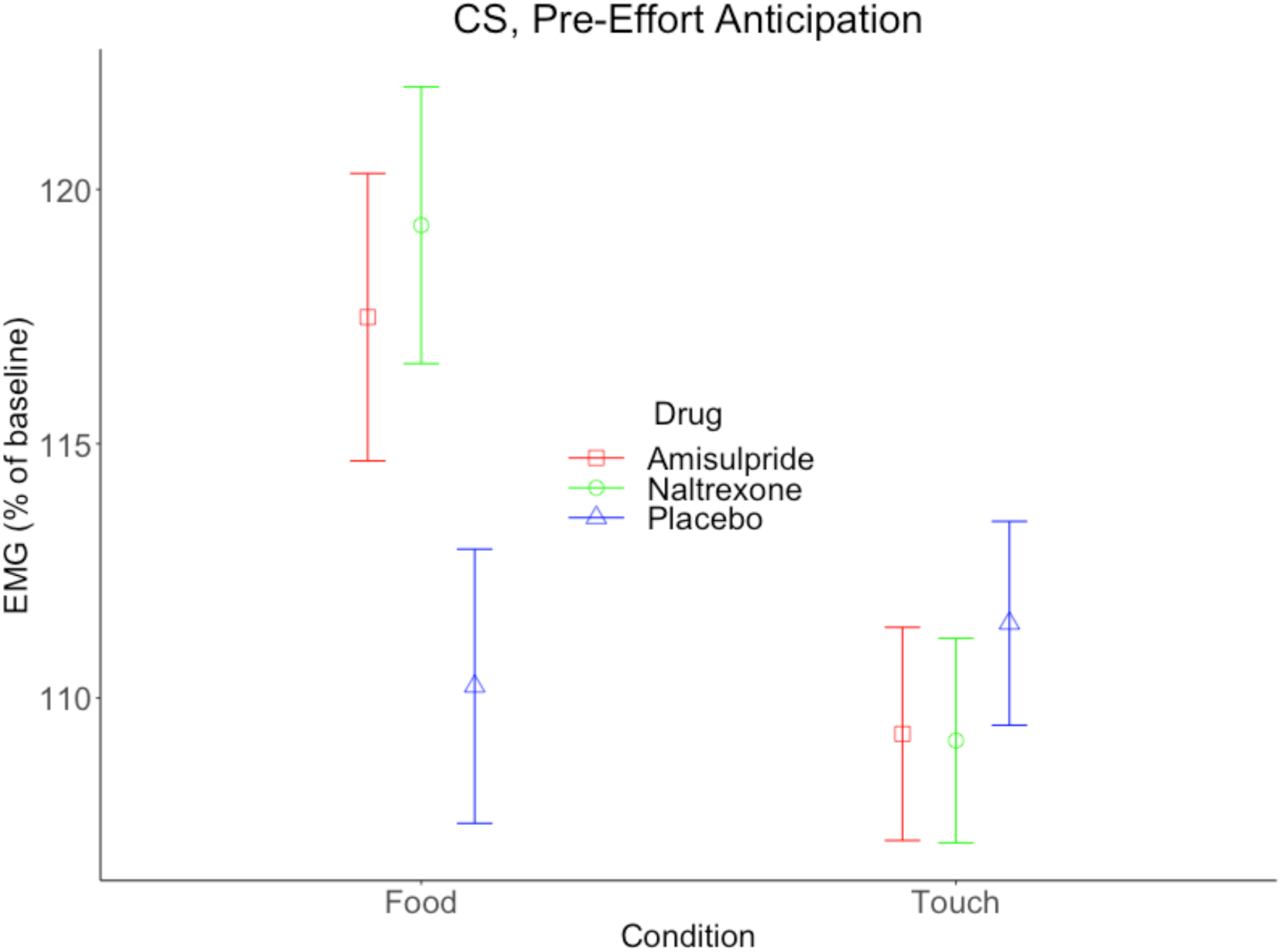
Marginal means (and 95% CIs) of the EMG of the CS muscle in the Pre-Effort anticipation window. A significant Condition x Drug interaction was found for the model including the predictor Wanting (shown here), and similarly for the model including the predictor Effort.

Similarly, a significant Drug X Condition interaction was found in the analysis of the CS muscle by Effort (*F*(2, 135.26) = 5.09, *p* = .007), reflecting greater CS activation to food than touch in the amisulpride (*p* = .002) and naltrexone group (*p* < .001), while the placebo group had similar activations across both conditions (*p* = .71). For the ZM muscle by Wanting and by Effort, no significant main effects or interactions involving the factor Drug were found (all *F* < 2, all *p* > .1).

In summary, activation of the CS in the Pre-Effort anticipation period was increased for food compared to touch stimuli in both active drug groups, but not in the placebo group, indicating more frowning when blocking the dopaminergic and the opioidergic systems.

### Post-Effort anticipation

No significant main or interaction effects involving the factor Drug were found for the CS nor the ZM muscle (all *F* < 1.9, all *p* > .15).

### Reward Delivery

No significant main or interaction effects involving the factor Drug were found for the CS by Liking (all *F* < 7, all *p* > .5). For the ZM, a significant Liking X Drug interaction was found (*F*(2, 128.80) = 3.97, *p* = .02). In the placebo group (Fig. 5), the slope for ZM activation with greater liking was significantly steeper than for the naltrexone group (*p* = .01) indicating impaired smiling during the delivery of liked rewards only when blocking the opioidergic system. Comparisons of placebo with amisulpride, and of amisulpride with naltrexone were not significant (all p > .2).

**Fig. 5:**
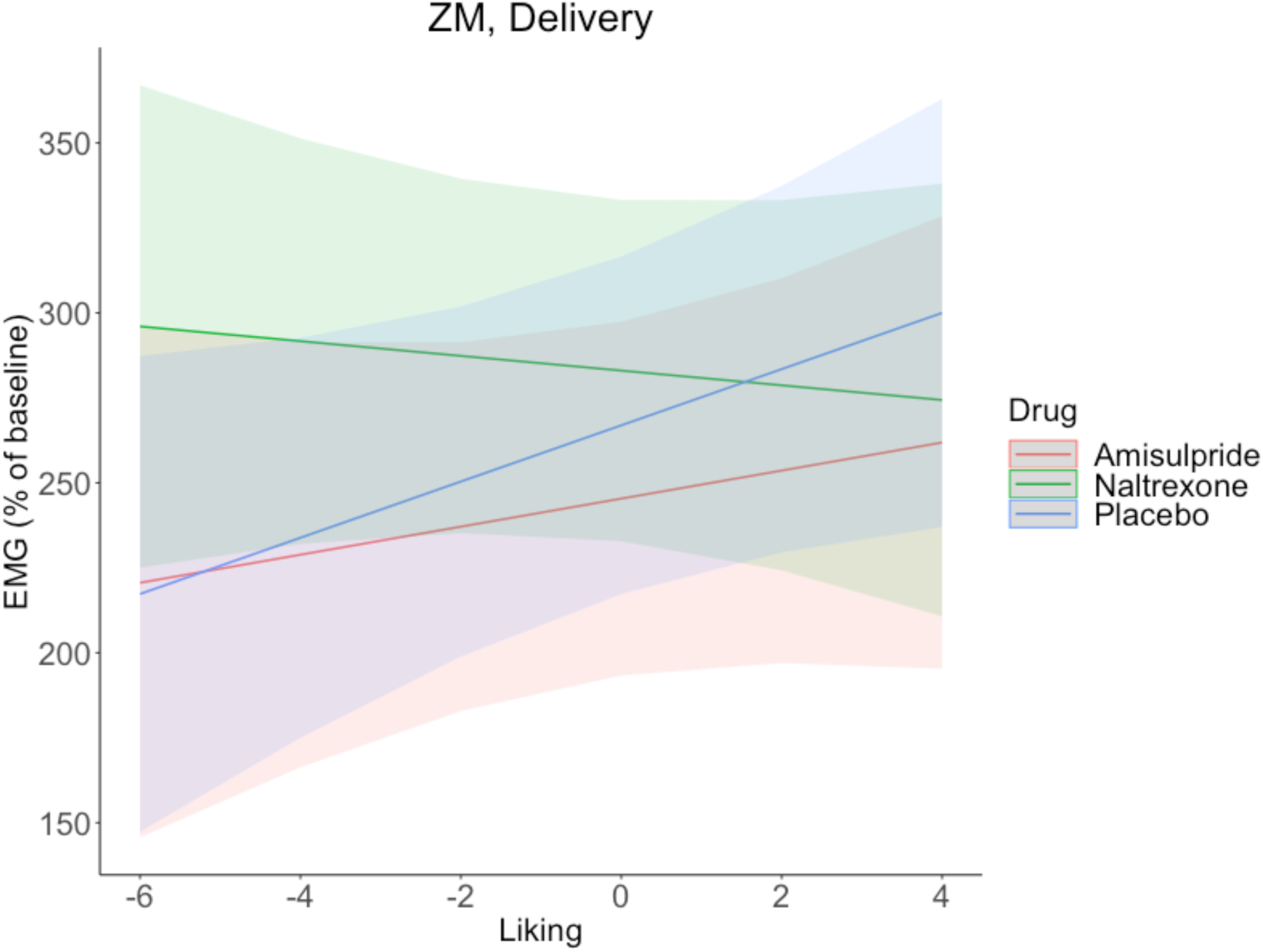
Model fit (and 95% CIs) of the ZM in the Delivery window. A significant Liking x Drug interaction reflected ZM activation for greater liking in the placebo group, but not in the naltrexone group, which showed the opposite pattern of ZM relaxation for greater liking.

### Relax phase

No significant main or interaction effects for the factor Drug were found in the CS and ZM data (all *F* < 1.7, all *p* > .19).

## Discussion

A recently developed experimental paradigm ^27^, in which reward processing is operationalised similarly to animal research, by assessing in each trial both explicit (ratings and effort) and implicit (facial EMG) anticipation and consumption of social and nonsocial rewards, was combined with a dopaminergic and opioidergic drug challenge. This allowed us to address two fundamental and as of yet unresolved research questions: 1) to which extent do motivational and hedonic responses in adult humans rely on shared or separate neurochemical systems, and 2) does the neurochemical basis of human reward processing differ for social and nonsocial rewards.

Analyses of the explicit measures ratings and effort revealed that participants who had taken either the D2/D3 dopamine receptor antagonist amisulpride, or the non-selective opioid receptor antagonist naltrexone, produced less effort compared to placebo to 1) obtain food rewards (Fig. 2A), and 2) to obtain low rewards of both conditions (Fig. 2B). These findings are in line with animal models indicating that both the dopaminergic and the opioidergic systems underlie the motivational aspect of reward processing ^14^. The fact that the effect was observed for the second-preferred (low) reward, but not for the most preferred (high) reward, rules out the possibility of a generic motor impairment (e.g. induced by dopaminergic blockage). Instead, the results speak for a genuine alteration of the incentive salience of the low reward, whose rewarding value approached that of the verylow reward, due to the pharmacological manipulation.

Interestingly, despite both drugs having an effect on the effort produced to obtain the low rewards, the amisulpride group also showed increased ratings of liking compared to the naltrexone group (Fig. 2C). This seems to corroborate the hypothesis that the dopaminergic system underlies the motivational but not the hedonic component of rewards, while the opioidergic system underlies both. However, these findings should be interpreted with caution, since liking of low rewards was also lower in the placebo compared to the amisulpride group.

To verify whether implicit facial reactions reflected the hedonic value of rewards, fEMG data was analyzed in a first step without the factor Drug. The findings confirmed the pattern expected based on prior work ^27,28,30,40,41^, of less frowning and more smiling during anticipation and consumption of most wanted and most liked rewards (Fig. 3). As we have recently reported using the same paradigm ^27^, most effects were in the CS muscle, which relaxed for greater wanting and effort in the Pre-Effort anticipation period, and for greater liking in the Delivery and Relax periods (the latter effect was stronger in the food condition). The ZM muscle showed the opposite pattern of activation for greater wanting in the Pre and Post-Effort anticipation periods, and for greater liking in the Delivery period (the first and last effect were only present in the food condition). The sensitivity of fEMG for capturing wanting and liking was thus confirmed, although the drug-independent effects were, as previously reported ^27^, more prominent for the CS than the ZM muscle.

In a second next step, we assessed the impact of the drugs on fEMG activity. In line with our initial hypothesis, only the opioidergic antagonist had an effect on implicit hedonic facial reactions during reward *consumption*. In particular, an increased ZM activation for greater liking was found in the placebo group during Delivery, and the slope of this effect was significantly different in the placebo than in the naltrexone group, which instead showed the opposite pattern of ZM relaxation with greater liking (Fig. 5). The fact that administration of the opioid antagonist naltrexone impaired smiling during the delivery of liked rewards parallels animal observation of reduced orofacial hedonic reactions to sweet food after opiodergic blockage^42^. Importantly, this finding cannot be explained by mouth movements that might have occurred during food delivery (although instruction to swallow followed the Delivery window), as statistics did not reveal an interaction with the factor Condition.

Confirming our hypothesis about implicit hedonic facial reactions during the *announcement* of a reward, both amisulpride and naltrexone resulted in greater CS activation during the Pre-Effort anticipation of food (Fig. 4). Because frowning typically reflects a more negative (or less positive) reaction^43,44^, the fact that dopamine and opioid antagonists led to greater frowning during reward anticipation might be interpreted as a reflection of less anticipated pleasure in these groups of participants.

Taken together, fEMG data showed a differential action of dopaminergic and opioidergic drug manipulation during the anticipation and consumption of rewards. In line with the explicit measures (ratings of wanting, ratings of liking and effort), the implicit measure fEMG indicates an effect of both amisulpride and naltrexone during the anticipation of rewards, but only of naltrexone during subsequent reward consumption. This pattern of results confirms and extends to human adults previous animal evidence about the role of the dopaminergic system for the motivational component and of the opioidergic system for both motivational and hedonic components of reward processing. This is the first evidence suggesting that the neurochemical regulation of pleasure (as indicated by hedonic facial reactions) is phase-specific, depending on whether the reward is anticipated or experienced.

Inclusion of both social (touch) and nonsocial (food) rewards allowed us to address the still unresolved question ^38^, whether different types of rewards are processed in the same neural structures (as proposed by the ‘common currency hypothesis’), or if representations coding for different rewards occur in distinct neural structures, albeit on a common scale ^45^. Social rewards in particular may constitute a separate class of stimuli, with a dedicated neural circuitry ^46^, which can be specifically impaired, for example in people with autism spectrum disorders ^47,48^. Although the magnitude of the two types of rewards was carefully matched^27^, we found that most drug effects were either stronger or restricted to food trials, e.g. in the Pre-Effort anticipation period. This suggests that wanting and liking of both social and nonsocial rewards may not rely on exactly the same neurochemical brain mechanisms. However, fEMG results to food were also stronger to begin with, which might explain why only this condition showed drug-induced effects. Another possible explanation for the less pronounced drug effects in the touch condition is that responses to social rewards, including touch, might also depend on oxytocin and serotonin, in addition to dopamine and opioids ^49,50^. Future studies should investigate the role of further neurochemical systems in the processing of social vs. nonsocial rewards. Ultimately, a clear answer to the question if different rewards are processed in the same or different brain areas may require the use of brain imaging, or of more direct measures of brain activity, in addition to pharmacological challenges tailored to investigate the role of different neurochemical systems in the processing of social vs. nonsocial rewards.

## Conclusion

We report pharmacological evidence in healthy human volunteers, across several measures including the monitoring of facial expressions with fEMG, for the hypothesis that *liking* of rewards relies only on the opioidergic system, and *wanting* of rewards relies on both the dopaminergic and opioidergic system. Interestingly, administration of both dopaminergic and opioidergic antagonists was accompanied by increased negative facial expressions during reward anticipation - suggesting that important neurochemical differences underlie hedonic expressions occurring during anticipation and experience of pleasure. This constitutes the first demonstration of this kind in adult humans, using an operationalization of reward closely resembling previous animal research. The finding that most drug effects were either stronger for or restricted to food trials potentially points to different neurochemical brain mechanisms for social and nonsocial rewards.

## Materials and Methods

### Subjects

Based on previous work that had used the same compounds and doses ^18^, we aimed at collecting data from 40 participants per group or more. The final participants sample included 131 volunteers (88 females) aged 18–35 years (*M* = 23.3; *SD* = 3.5). In the amisulpride group blood concentrations of the drug (measured five hours after intake) were in or above the therapeutic range (blood samples missing for six people). Specifically, the minimum was 212 ng/ml, and 19 participants were above 604 ng/ml. All participants reported being right-handed, to smoke less than five cigarettes daily, to have no history of current or former drug abuse, to like milk and chocolate, not to suffer from diabetes, lactose intolerance, lesions or skin disease on the left forearm, and to be free of psychiatric or neurological disorders. Participants’ average Body Mass Index (BMI) was 22.6, (*SD* = 2.5, range 17.7 – 29.3). To reduce the chances that social touch would be perceived as a sexual reward, the social touch stimulation was always carried out by a same-sex experimenter (see Procedure), and only participants who reported to be heterosexual were included. The study was approved by the Ethical Committee of the Medical University of Vienna (EK N. 1393/2017) and was performed in line with the Declaration of Helsinki ^51^. Participants signed informed consent and received a monetary compensation of 90€.

### Stimuli

Three stimuli with identical fat and sugar content (1.5 g fat, 10 g of sugar per 100 g) were used as rewards in the Nonsocial condition: milk, chocolate milk, and a 4:1 mix of milk and chocolate milk. Tap water served for rinsing at the end of each trial. The initial stimulus temperature of these liquids was kept constant (∼4° C) across participants. Stimulus delivery was accomplished through computer-controlled pumps (PHD Ultra pumps, Harvard Apparatus) attached to plastic tubes (internal ø 1,6 mm; external ø 3,2 mm; Tygon tubing, U.S. Plastic Corp.), which ended jointly on an adjustable mount positioned about two centimeters in front of the participant’s mouth. In each trial, two ml of liquid were administered during two seconds. Overall, including stimulus pretesting (see Procedure), participants consumed 196 ml of liquids, composed of 98 ml of water, and 98 ml of sweet milk with different concentrations of chocolate aroma (depending on effort, see below).

Social rewards consisted of gentle caresses over a previously-marked nine-cm area of the participant’s forearm (measurement started from the wrist towards the elbow). Three different caressing frequencies, chosen based on the literature and pilot testing, were applied during six seconds by a same-sex experimenter: six cm/s, 21 cm/s and 27 cm/s. To facilitate stroking, the stimulating experimenter received extensive training and in each trial heard rhythmic sounds, indicating the rhythm for stimulation, through headphones.

### EMG

After cleansing of the corresponding face areas with alcohol, water, and an abrasive paste, reusable Ag/AgCl electrodes with 4 mm inner and 8 mm outer diameter were attached bipolarly according to guidelines ^52^ on the left corrugator supercilii (CS) and the zygomaticus major (ZM) muscles. A ground electrode was attached to the participants’ forehead, and a reference electrode on the left mastoid. EMG data was sampled at 1200 Hz with impedances below 20 kOHM using a g.USBamp amplifier (g.tec Medical Engineering GmbH) and the software Matlab (MathWorks, Inc.).

### Procedure

A monocentric, randomized, double-blind, placebo-controlled, three-armed study design was used. The study took place in the Department of Psychiatry and Psychotherapy at the Medical University of Vienna. Participants visited the laboratory for a first visit (T0) in which they received a health screening, followed by a second visit (T1) that included oral drug intake and the experiment described here. Pharmacological dosage, and length of waiting time after drug intake (three hours) were modeled on previous work ^18^.

Participants came to T1 with an empty stomach (they were instructed not to eat in the preceding six hours), filled out the PANAS questionnaire, tested negative on a urine drug screen sensitive to opiates, amphetamine, methamphetamine, cocaine (among other things), and then received a capsule filled with either 400 mg of amisulpride (Solian®), 50 mg of naltrexone (Dependex®), or 650 mg of mannitol (sugar) from the study doctor. All capsules looked identical from the outside, and neither participants nor the experimenters were informed of their content. Drug intake was followed by a waiting period, EMG preparation, and task instructions.

The experiment comprised two tasks following procedures described elsewhere ^27^. The main task started three hours after pill intake. Participants were seated at a table and comfortably rested their left forearm on a pillow. A curtain blocked their view of the left forearm and the rest of the room. This was particularly relevant for the social condition, in which one of two same-sex experimenters applied the social rewards to the participant’s left forearm. Two experimenters were always present during testing, to limit the influence of participants’ experimenter preferences, and to allow participants to better concentrate on the (social) stimuli.

Participants first completed a short task in which they experienced and individually ranked three reward levels with liking-ratings for the social and nonsocial condition and the respective stimuli, presented randomly in sets of threeof the same condition. In the main task, which started three hours after pill intake, the previously most liked stimuli were used as ‘high’ rewards, the stimuli with medium liking as ‘low’ rewards, and the least liked stimuli were used as ‘verylow’ rewards. To calibrate the dynamometer, the maximum voluntary contraction (MVC) was established right before the short task, by asking participants to squeeze the dynamometer (HD-BTA, Vernier Software & Technology, USA) with their right hand as hard as possible three times during three seconds. The average MVC (peak force in newtons across all three trials) was 212 (*SD* = 80.4), and did not differ between Drug groups, as tested by linear regression (*β* = 1.6, *SE* = 8.68, *t* = 0.19, *p* = 0.85).

After calibration of the dynamometer, EMG electrodes were attached, participants received detailed instructions, and completed four practice trials (two per condition). The main task included four experimental blocks with 20 trials each. Each block contained eitherfood or touch trials, and the blocks were interleaved (ABAB or BABA) in a counterbalanced order across participants. Each trial included (Fig. 1) the following main steps (see Supporting Information for all elements of a trial): 1) a picture announcing the highest possible reward (high or low, 3 sec), 2) a continuous scale ranging from ‘not at all’ to ‘very much’ to rate (without a time limit) wanting of the announced reward (ratings were converted to a Likert scale ranging from −10 to +10), 3) a 4-sec period of physical effort, during which probability of receiving the announced reward was determined by the amount of force exerted by squeezing the dynamometer with the right hand, while receiving visual feedback (sliding average of 1 sec, as percentage of the MVC), 4) a picture announcing the obtained reward (3 sec in the nonsocial, 7.3 sec in the social condition), which could be high, low, or – if insufficient effort had been exerted – verylow (the greater participants’ effort, the higher the probability of obtaining the announced reward), 5) a phase of reward delivery (2 sec in the food, 6.5 sec in the touch condition – this difference in timing was necessary to obtain sufficiently long tactile stimulation, while keeping the overall trial duration similar across conditions), 6) in the food condition instructions to lean back and swallow the obtained reward (duration 3 sec), 7) a relaxation phase (5 sec), and 8) a continuous scale to rate the liking of the obtained reward. In the food condition, participants then received water for mouth rinsing. In both conditions trials ended with a blank screen for 3 to 4 seconds. The last four trials in each block did not require pressing of the dynamometer – these trials were kept in the data, as removing them from analyses did not change the pattern of results. After each block participants were allowed to take a short break.

Both tasks were run on a desktop computer with Windows 7 using MATLAB 2014b and the Cogent 2000 and Cogent Graphics toolboxes, and presented on an LCD monitor with a resolution of 1280 x 1024 pixels. The Positive and Negative Affect Schedule (PANAS) ^39^ was filled out twice at the main laboratory visit: just before pill intake, and 3 hours later. Levels of amisulpride (ng/ml) were measured in blood samples taken five hours after pill intake (after both tasks).

### Analyses

Data were analyzed with linear mixed effects models (LMMs) using the lmer() function of the *lme4* package in R ^53,54^ (with exception of group comparisons for age, BMI, MVC, and PANAS scores, and number of excluded trials, which were made with linear regressions using the lm() function). In comparison to ANOVAs, LMMs reduce Type-I errors and allow for the generalization of findings ^55^. All the data and analysis scripts are available online (https://bit.ly/35UtUvw). Figures (except Fig. 1) were created in R using the packages *ggplot2, ggpirate,* and *cowplot*.

Behavioral data were analyzed in the following manner. Outlier trials were defined as those with a rating of wanting, rating of liking, or amount of exerted force, which was greater/smaller than the subject’s mean +/− 2 times the subject’s standard deviation. This led to an average rejection of 6.56 trials per participant (*SD* = 3.71). The number of excluded trials did not differ between groups, as tested by linear model (*F*(2, 128) = 2.54, *p* = 0.08). For each behavioral dependent variable (ratings of wanting and liking, effort), a LMM was run with the fixed effects Condition (food, touch), RewardType (high, low, verylow), and Drug (amisulpride, naltrexone, placebo). Categorical predictors were centered through effect coding, and by-subject random intercepts and slopes for all within-subjects factors and their interactions were included as random effects (unless the model did not converge, in which case first the interaction among within-subjects factors and then the slopes by Condition were dropped). Type-III F-tests were computed with the Satterthwaite degrees of freedom approximation, using the anova() function of the *lmerTest* package.

Due to technical failure, one participant lacked EMG data entirely, and another participant lacked EMG for half of the trials. EMG data were preprocessed in Matlab R2018a (www.themathworks.com), partly using the EEGLAB toolbox ^56^. A 20 to 400 Hz bandpass filter was applied, then data were rectified and smoothed with a 40 Hz low-pass filter. Epochs were extracted focusing on periods of reward anticipation (Pre-Effort and Post-Effort anticipation) and reward consumption (Delivery and subsequent Relax). EMG was averaged over time-windows of one second, with exception of the 6.5-seconds-long period of touch Delivery, which was averaged over five windows of 1.3 seconds each, to obtain the same number of windows as for the food condition. We excluded for each participant trials on which the average amplitude in the baseline period (one second during fixation) of the CS or ZM muscles was lower than M−2*SD, or higher than M+2*SD (M = average amplitude over all trials’ baselines for the respective muscle and participant). On average, this led to the rejection of 7.7 % of trials per participant (*SD* = 2.5). EMG analyses were carried out in four periods of interest: *Pre-effort anticipation* during reward announcement at the beginning of each trial (3 sec), *Post-effort anticipation* during the announcement of the gained reward (3 sec), *Delivery* (5 sec in the Food and 6.5 sec in the Touch condition, both averaged to five 1− sec time windows), and *Relax* (5 sec). For each trial, values in these epochs were expressed as percentage of the average amplitude during the fixation cross at the beginning of that trial. For the Pre- and Post-Effort anticipation periods, separate linear mixed-effects models (LMM) were fitted by muscle, with the fixed effects Drug (amisulpride, naltrexone, placebo), Condition (food, touch), and either trial-by-trial Wanting, or Effort (continuous predictors). During the Post-Effort anticipation period participants could receive the information that they were going to obtain the verylow reward, to which the preceding ratings of wanting and effort did not apply. Therefore, verylow trials were excluded from analyses of the Post-Effort anticipation period by Wanting and Reward. For the Delivery and Relax periods, separate LMMs on all trials were fitted by muscle, with the fixed effects Drug, Condition, and Liking. In all LMMs Wanting, Effort, and Liking were centered and scaled by subject.

## Acknowledgments

The study was supported by the Vienna Science and Technology Fund (WWTF) with a grant (CS15-003) awarded to Giorgia Silani and Christoph Eisenegger, and a grant (VRG13-007) awarded to Christoph Eisenegger. Funders had no role in study design, data collection and analysis, decision to publish or preparation of the manuscript. We thank Prof. Boris Quednow for commenting on an earlier version of the manuscript. Many thanks to Mumna Al Banchaabouchi and the interns and master students whose help was crucial for data acquisition: Mani Erfanian Abdoust, Anne Franziska Braun, Raimund Bühler, Lena Drost, Manuel Czornik, Lisa Hollerith, Berit Hansen, Luise Huybrechts, Merit Pruin, Vera Ritter, Conrad Seewald, Frederic Schwetz, Carolin Waleew, Luca Wiltgen, Stephan Zillmer.

## Author contributions

S.K., G.S., C.E conception and design of the work. S.G. supervised data collection. P.S., I.G., and M.W. were responsible for all medical aspects and for drug delivery. S.K. performed data processing and analysis, to which G.S. and C.M. provided valuable input. S.K. and G.S. wrote the manuscript, with input from all co-authors.

## Supporting Information

### 1. Description of all elements in a trial

Each trial included (Fig. S1) the following steps: 1) a fixation cross (2 sec), 2) a picture announcing the highest possible reward (high or low, 3 sec), 3) a continuous scale ranging from ‘not at all’ to ‘very much’ to rate (without time limit) wanting of the announced reward (ratings were converted to a 20-point Likert scale), 4) a 2-sec period announcing the effort phase, 5) a 4-sec period of physical effort, during which participants could set the chances of receiving the announced reward by the amount of force they would exert by squeezing the dynamometer with their right hand while receiving visual feedback of the amount of exerted force (sliding average of 1 sec, as percentage of the MVC), 6) a picture announcing the obtained reward (3 sec), which could be high, low, or verylow (the greater participants’ effort, the higher the probability of obtaining the announced reward), 7) a phase of preparation for reward delivery (3 sec in the food, 7.3 sec in the touch condition), 8) a phase of reward delivery (2 sec in the food, 6.5 sec in the touch condition – this difference in timing was necessary to obtain sufficiently long tactile stimulation, while keeping the overall trial duration similar across conditions), and only in the food condition also instructions to slightly lean back and swallow the obtained reward (duration 3 sec), 9) a relaxation phase (5 sec), and 10) a continuous scale ranging from negative to positive to rate the liking of the obtained reward. In the food condition, participants then received water for mouth rinsing, in a way similar to how they had received the food reward. In both conditions trials ended with a blank screen for 3 to 4 seconds.

**Fig. S1:**
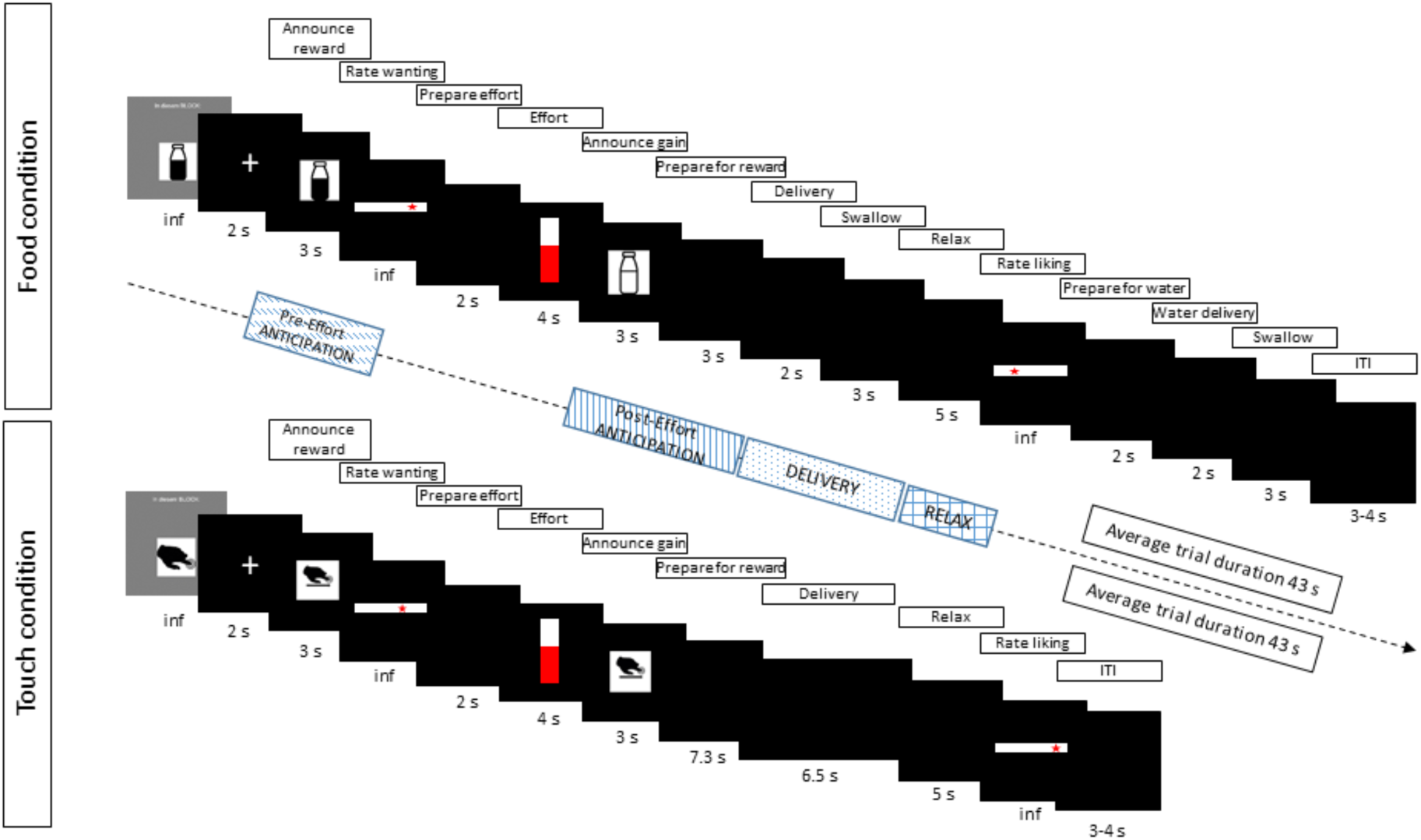
Trial examples for the food and touch conditions, including all screens.

### 2. Full statistical results

#### Explicit measures of Social and Nonsocial reward: wanting, liking and effort

The number of trials with high, low, or verylow rewards did not differ across groups, as shown by a linear mixed effects model (LMM) with number of trials as dependent variable, the fixed effects Condition (food, touch), Drug (amisulpride, naltrexone, placebo), and RewardType (high, low, verylow), and by-subject random intercepts. Only a significant main effect of Reward Type was found (*F*(2, 763) = 27.84, *p* < .001), due to a greater number of high (*M* = 16.53, *SD* = 2.87) than low (*M* = 14.93, *SD* = 3.46) and verylow (*M* = 8.67, *SD* = 4.93) trials across all three groups.

Moreover, groups of participants did not differ in their maximum voluntary contraction (MVC) of the hand dynamometer, which was measured right before the main task (*β* = 1.6, *SE* = 8.68, *t* = 0.19, *p* = 0.85), nor in their positive and negative mood measured with the PANAS at time of pill intake or three hours later (all *β* < 0.6, all *t* < 0.8, all *p* > 0.4).

Subjective ratings of wanting and liking, and effort, were analyzed in separate linear mixed effects models (LMMs) with Condition (food, touch), Drug (amisulpride, naltrexone, placebo), and RewardType (high, low announced at the beginning of each trial for wanting and effort; high, low, verylow obtained after the effort phase for liking) as fixed effects, and as random effects intercepts for subjects and by-subject random slopes for the effects of Condition, RewardType, and their interaction.

**Fig. S2:**
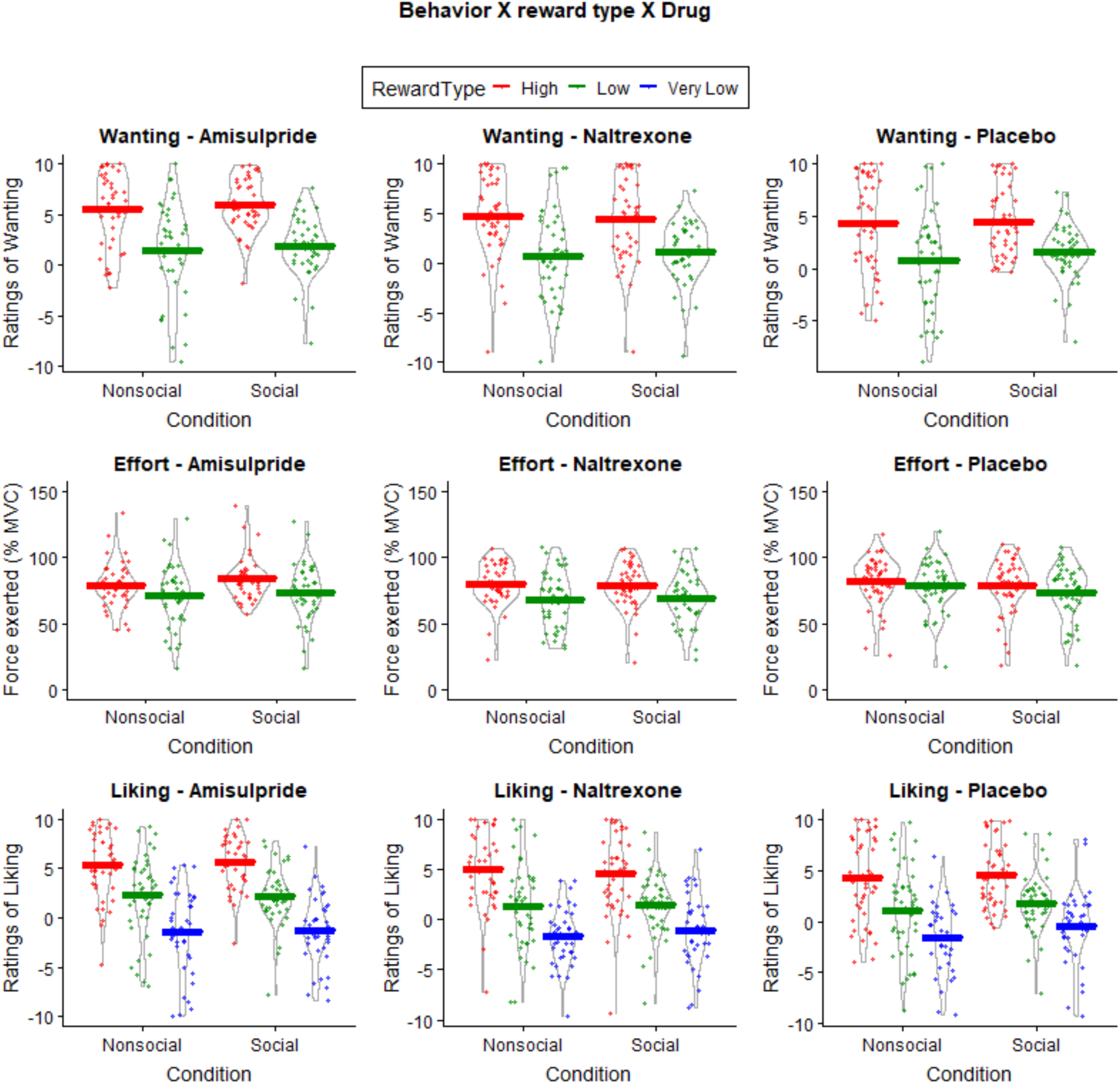
Violin plots depicting ratings of wanting (top row), physical effort (middle row), and ratings of liking (bottom row) for nonsocial (food) and social (touch) rewards, divided by drug group. Ratings of wanting and liking were recorded on Likert scales ranging from −10 (‘not at all’) to +10 (‘very much’). Exerted force is the maximum value (as percentage of the MVC) reached in a 4-sec period. These measures are shown as a function of RewardType, i.e. the individual preferences of three social and three nonsocial stimuli, measured at the beginning of the experiment. Each data point corresponds to the mean per participant, horizontal lines show means over all participants, outer lines represent the probability densities. The verylow rewards were never announced, and were only received after insufficient effort had been exerted – they therefore only appear in the ratings of liking.

Behavioral analyses on ratings of wanting (Fig. S2, top) resulted in an expected significant main effect of RewardType (*F*(1, 128.02) = 119.3159, p < .001), due to higher ratings of wanting for high reward (*M* = 4.89, *SD* = 4.31) compared to low reward (*M* = 1.14, *SD* = 4.46). All other effects were not significant (all *F* < 2.4, all *p* > .1).

The same LMM on effort (Fig. 2, center) resulted in the expected significant main effect of RewardType (*F*(1, 128.49) = 54.1821, *p* < .001), due to stronger force applied for high (*M* = 80.49, *SD* = 22.35) than low rewards (*M* = 71.74, *SD* = 25.42); a significant Condition X Drug interaction (*F*(1, 128.31) = 4.5369, *p* = .01) reflecting lower effort in the food condition in the amisulpride (*M* = 74.98, *SD* = 26.57) and naltrexone (*M* = 73.51, *SD* = 24.43) groups compared to the placebo (*M* = 80.20, *SD* = 22.41) group, but similar force across drug groups in the Social condition (amisulpride: *M* = 78.34, *SD* = 25.14; naltrexone: *M* = 73.78, *SD* = 23.15; placebo: *M* = 76.11, *SD* = 23.51). A marginally significant RewardType X Drug interaction was also found (*F*(1, 128.50) = 2.9734, *p* = .055), due to reduced effort to low rewards in the amisulpride (*M* = 71.67, *SD* = 27.60) and naltrexone (*M* = 67.90, *SD* = 24.45) groups compared to the placebo group (*M* = 75.65, *SD* = 23.60), but no differences in effort between groups for high rewards (amisulpride: *M* = 81.60, *SD* = 23.09; naltrexone: *M* = 79.29, *SD* = 21.70; placebo: *M* = 80.63, *SD* = 22.23). All other effects were not significant (all *F* < 0.9, all *p* > .4).

The same LMM on the liking ratings (Fig. 2, bottom) resulted in the expected main effect of RewardType (*F*(2, 260.22) = 116.0760, *p* < .001), with greatest liking of high rewards (*M* = 5.20, *SD* = 3.92), followed by low rewards (*M* = 2.00, *SD* = 4.06), and verylow rewards at the bottom (*M* = −1.26, *SD* = 3.95; all pairwise comparisons *p* < .001). A significant Drug X RewardType interaction (*F*(4, 260.24) = 8.2899, *p* < .001) reflected greater liking of low rewards in the amisulpride group (*M* = 2.57, *SD* = 3.78) compared with both the naltrexone (*M* = 1.33, *SD* = 4.35, *p* = .01, *p* = .01) and the placebo groups (*M* = 1.53, *SD* = 4.18, *p* = .02, *p* = .02).

In summary, main effects of RewardType were found across all behavioral measures, reflecting greater wanting and liking of high compared to low rewards. The amisulpride and naltrexone groups showed reduced effort to obtain food rewards, and to obtain low rewards of both conditions – although these effects did not survive post-hoc pairwise comparisons. The amisulpride group also showed greater liking of low rewards compared to both the naltrexone and placebo groups.

#### Implicit measures of food and touch rewards: facial EMG

##### Pre-Effort-Anticipation

For the CS muscle by Wanting, significant main effects of Condition (*F*(1, 128.73) = 12.0021, *p* < .001) and Wanting (*F*(1, 164.72) = 10.6538, *p* = .001) were found. Activation of the CS was greater for the food (*M* = 116.35, *SD* = 112.85) than the touch (*M* = 110.21, *SD* = 81.59) condition and decreased, as expected, with increasing ratings of wanting (b = −6.5). A significant Drug X Condition interaction (*F*(2, 128.70) = 4.8080, *p* = .009) reflected greater CS activation in to food than touch in the amisulpride group (*p* = .004; food: *M* = 119.12, *SD* = 134.43; touch: *M* = 109.46, *SD* = 76.09) and naltrexone group (*p* < .001; food: *M* = 120.00, *SD* = 128.19; touch: *M* = 109.66, *SD* = 89.46), while the placebo group had similar activations across both conditions (*p* = .68; food: *M* = 110.30, *SD* = 65.72; touch: *M* = 111.44, *SD* = 78.17). All other effects were not significant (all *F* < 2.2, all *p* > .1). CS activation in the food condition was also significantly greater in the naltrexone than placebo group (*p* = .04).

For the CS muscle by Effort, in addition to the aforementioned effects of Condition and Drug X Condition, a significant main effect of Effort was found (*F*(1, 148.37) = 10.0506, *p* = .002), due to CS relaxation with increasing levels of Effort (b = - 6.04).

For the ZM muscle by Wanting, a significant main effect of Condition (*F*(1, 128.12) = 13.9723, *p* < .001) was also found, reflecting greater ZM activation for food (*M* = 138.79, *SD* = 185.47) than touch (*M* = 122.98, *SD* = 168.20) rewards. Moreover, a significant Condition X Wanting interaction (*F*(1, 122.00) = 6.0828, *p* = .02) reflected increased ZM activation with increasing ratings of Wanting in the food (b = 5.25, *p* = .02) but not in the Touch condition (b = −3.81, p = .34). All other effects were not significant (all *F* < .7, all *p* > .54).

For the ZM muscle by Effort (random slopes for the Condition X Effort interaction were removed to allow model convergence), only a significant main effect of Condition was found (*F*(1, 127.4) = 14.6601, *p* < .001), with greater ZM activation during food (*M* = 138.79, *SD* = 185.47) compared to touch (*M* = 122.98, *SD* = 168.20) trials.

In summary, activation of the CS in the Pre-Effort anticipation period was inversely related to Wanting and Effort, and was increased for food compared to touch stimuli in both active drug groups, but not in the placebo group. In the food condition, greater ZM activation for increasing Wanting was also found.

##### Post-Effort anticipation

No significant effects were found for the CS muscle, neither by Wanting, nor by Effort (all *F* < 2.3, all *p* > .1).

For the Zygomaticus, only a significant main effect of Wanting was found (*F*(1, 130.33) = 6.5565, *p* = .01), due to greater ZM contraction for increasing levels of Wanting (b = −6.67).

##### Reward Delivery

Analysis of the CS resulted in a significant main effect of Condition (*F*(1, 125.16) = 5.7899, *p* = .02), due to greater muscle activation in the food (*M* = 150.44, *SD* = 202.56) than touch condition (*M* = 116.79, *SD* = 395.75), and in a trend for a main effect of Liking (*F*(1, 113.08) = 3.2671, *p* = .07).

For the ZM a significant main effect of Condition (*F*(1, 126.79) = 74.5420, *p* < .001), as well as a significant Liking X Drug interaction (*F*(2, 125.21) = 3.2858, *p* = .04) were found. In the placebo group, the slope for ZM activation for greater liking was significantly steeper than for the naltrexone group (*p* = .03). No difference between placebo and amisulpride and between amisulpride and naltrexone groups emerged (all p > .3).

##### Relax phase

For the CS by Liking (only the random slope for Liking was included to allow model convergence), significant main effects of Condition (*F*(1, 22925.6) = 132.9776, *p* < .001) and Liking (*F*(1, 128.6) = 15.3266, *p* < .001), and significant Condition X Liking (*F*(1, 16300.1) = 6.3334, *p* = .01) and Condition X Drug (*F*(1, 22902.1) =3.7972, *p* = .02) interactions were found. The Condition X Liking interaction was due to the fact that while CS activation decreased with greater liking in general (b = −16.8), this effect was stronger in the Food than the Touch condition (*p* = .04). The Condition X Drug interaction was due to greater CS activation to food rewards in the naltrexone group (*M* = 161.22, *SD* = 479.78) than in the amisulpride (*M* = 132.93, *SD* = 191.94; *p* = .04) and placebo (*M* = 134.36, *SD* = 138.97; *p* = .05) groups. CS activations did not differ between groups in the touch condition (all *p* > .6).

For the ZM, only a significant main effect of Condition was found (*F*(1, 127.17) = 127.5590, *p* < .001), reflecting greater ZM contraction in the food (*M* = 211.18, *SD* = 218.25) than in the touch (*M* = 133.32, *SD* = 289.47) condition.

## References

1. Schultz, W. Reward. in Brain Mapping: An Encyclopedic Reference (ed. Toga, A. W.) vol. 2 643–651 (Academic Press: Elsevier, 2015).

2. Berridge, K. C. Food reward: brain substrates of wanting and liking. Neurosci. Biobehav. Rev. 20, 1–25 (1996).

3. Berridge, K. C. & Kringelbach, M. L. Pleasure systems in the brain. Neuron 86, 646–664 (2015).

4. Berridge, K. C. & Robinson, T. E. What is the role of dopamine in reward: hedonic impact, reward learning, or incentive salience? Brain Res. Brain Res. Rev. 28, 309–369 (1998).

5. Kahneman, D., Wakker, P. P. & Sarin, R. Back to Bentham? Explorations of Experienced Utility. Q. J. Econ. 112, 375–406 (1997).

6. Berridge, K. C. & O’Doherty, J. P. From Experienced Utility to Decision Utility. in Neuroeconomics (Second Edition) (eds. Glimcher, P. W. & Fehr, E.) 335–351 (Academic Press, 2014). doi:10.1016/B978-0-12-416008-8.00018-8.

7. Berridge, K. C. Evolving Concepts of Emotion and Motivation. Front. Psychol. 9, (2018).

8. Der-Avakian, A., Barnes, S. A., Markou, A. & Pizzagalli, D. A. Translational Assessment of Reward and Motivational Deficits in Psychiatric Disorders. Curr. Top. Behav. Neurosci. 28, 231–262 (2016).

9. Grill, H. J. & Norgren, R. The taste reactivity test. I. Mimetic responses to gustatory stimuli in neurologically normal rats. Brain Res. 143, 263–279 (1978).

10. Barbano, M. F. & Cador, M. Opioids for hedonic experience and dopamine to get ready for it. Psychopharmacology (Berl.) 191, 497–506 (2007).

11. Berridge, K. C., Venier, I. L. & Robinson, T. E. Taste reactivity analysis of 6-hydroxydopamine-induced aphagia: implications for arousal and anhedonia hypotheses of dopamine function. Behav. Neurosci. 103, 36–45 (1989).

12. Treit, D. & Berridge, K. C. A comparison of benzodiazepine, serotonin, and dopamine agents in the taste-reactivity paradigm. Pharmacol. Biochem. Behav. 37, 451–456 (1990).

13. Berridge, K. C. & Valenstein, E. S. What psychological process mediates feeding evoked by electrical stimulation of the lateral hypothalamus? Behav. Neurosci. 105, 3–14 (1991).

14. Peciña, S. & Smith, K. S. Hedonic and motivational roles of opioids in food reward: implications for overeating disorders. Pharmacol. Biochem. Behav. 97, 34–46 (2010).

15. Taha, S. A. Preference or fat? Revisiting opioid effects on food intake. Physiol. Behav. 100, 429–437 (2010).

16. Callesen, M. B., Scheel-Krüger, J., Kringelbach, M. L. & Møller, A. A systematic review of impulse control disorders in Parkinson’s disease. J. Park. Dis. 3, 105–138 (2013).

17. Evans, A. H. et al. Compulsive drug use linked to sensitized ventral striatal dopamine transmission. Ann. Neurol. 59, 852–858 (2006).

18. Weber, S. C. et al. Dopamine D2/3- and μ-opioid receptor antagonists reduce cue-induced responding and reward impulsivity in humans. Transl. Psychiatry 6, e850 (2016).

19. Soutschek, A. et al. The dopaminergic reward system underpins gender differences in social preferences. *Nat*. Hum. Behav. 1 (2017) doi:10.1038/s41562-017-0226-y.

20. Eikemo, M. et al. Sweet taste pleasantness is modulated by morphine and naltrexone. Psychopharmacology (Berl.) 233, 3711–3723 (2016).

21. Chelnokova, O. et al. Rewards of beauty: the opioid system mediates social motivation in humans. Mol. Psychiatry 19, 746–747 (2014).

22. Büchel, C., Miedl, S. & Sprenger, C. Hedonic processing in humans is mediated by an opioidergic mechanism in a mesocorticolimbic system. eLife 7, e39648 (2018).

23. Pool, E., Sennwald, V., Delplanque, S., Brosch, T. & Sander, D. Measuring wanting and liking from animals to humans: A systematic review. Neurosci. Biobehav. Rev. 63, 124–142 (2016).

24. Berridge, K. C. Measuring hedonic impact in animals and infants: microstructure of affective taste reactivity patterns. Neurosci. Biobehav. Rev. 24, 173–198 (2000).

25. Steiner, J. E., Glaser, D., Hawilo, M. E. & Berridge, K. C. Comparative expression of hedonic impact: affective reactions to taste by human infants and other primates. Neurosci. Biobehav. Rev. 25, 53–74 (2001).

26. Delamater, A. R., LoLordo, V. M. & Berridge, K. C. Control of fluid palatability by exteroceptive Pavlovian signals. J. Exp. Psychol. Anim. Behav. Process. 12, 143–152 (1986).

27. Korb, S. et al. Facial responses of adult humans during the anticipation and consumption of touch and food rewards. Cognition 194, 104044 (2020).

28. Mayo, L. M., Lindé, J., Olausson, H., Heilig, M. & Morrison, I. Putting a good face on touch: Facial expression reflects the affective valence of caress-like touch across modalities. Biol. Psychol. 137, 83–90 (2018).

29. Pawling, R., Cannon, P. R., McGlone, F. P. & Walker, S. C. C-tactile afferent stimulating touch carries a positive affective value. PloS One 12, e0173457 (2017).

30. Ree, A., Mayo, L. M., Leknes, S. & Sailer, U. Touch targeting C-tactile afferent fibers has a unique physiological pattern: A combined electrodermal and facial electromyography study. Biol. Psychol. 140, 55–63 (2019).

31. Franzen, J. & Brinkmann, K. Wanting and liking in dysphoria: Cardiovascular and facial EMG responses during incentive processing. Biol. Psychol. 121, 19–29 (2016).

32. Löken, L. S., Wessberg, J., Morrison, I., McGlone, F. & Olausson, H. Coding of pleasant touch by unmyelinated afferents in humans. Nat. Neurosci. 12, 547–548 (2009).

33. McGlone, F., Wessberg, J. & Olausson, H. Discriminative and Affective Touch: Sensing and Feeling. Neuron 82, 737–755 (2014).

34. Ackerley, R., Saar, K., McGlone, F. & Backlund Wasling, H. Quantifying the sensory and emotional perception of touch: differences between glabrous and hairy skin. Front. Behav. Neurosci. 8, 34 (2014).

35. Salamone, J. D. et al. Haloperidol and nucleus accumbens dopamine depletion suppress lever pressing for food but increase free food consumption in a novel food choice procedure. Psychopharmacology (Berl.) 104, 515–521 (1991).

36. Lopez-Persem, A., Rigoux, L., Bourgeois-Gironde, S., Daunizeau, J. & Pessiglione, M. Choose, rate or squeeze: Comparison of economic value functions elicited by different behavioral tasks. PLoS Comput. Biol. 13, e1005848 (2017).

37. Treadway, M. T., Buckholtz, J. W., Schwartzman, A. N., Lambert, W. E. & Zald, D. H. Worth the ‘EEfRT’? The Effort Expenditure for Rewards Task as an Objective Measure of Motivation and Anhedonia. PLOS ONE 4, e6598 (2009).

38. Ruff, C. C. & Fehr, E. The neurobiology of rewards and values in social decision making. Nat. Rev. Neurosci. 15, 549–562 (2014).

39. Watson, D., Clark, L. A. & Tellegen, A. Development and validation of brief measures of positive and negative affect: the PANAS scales. J Soc Psychol 54, 1063–70 (1988).

40. Gilbert, A. N., Fridlund, A. J. & Sabini, J. Hedonic and social determinants of facial displays to odors. Chem. Senses 12, 355–363 (1987).

41. Horio, T. EMG Activities of Facial and Chewing Muscles of Human Adults in Response to Taste Stimuli. Percept. Mot. Skills 97, 289–298 (2003).

42. Smith, K. S. & Berridge, K. C. Opioid Limbic Circuit for Reward: Interaction between Hedonic Hotspots of Nucleus Accumbens and Ventral Pallidum. J. Neurosci. 27, 1594–1605 (2007).

43. Heller, A. S., Greischar, L. L., Honor, A., Anderle, M. J. & Davidson, R. J. Simultaneous acquisition of corrugator electromyography and functional magnetic resonance imaging: a new method for objectively measuring affect and neural activity concurrently. NeuroImage 58, 930–934 (2011).

44. The Science of Facial Expression. (Oxford University Press, 2017).

45. Grabenhorst, F. & Rolls, E. T. Value, pleasure and choice in the ventral prefrontal cortex. Trends Cogn. Sci. 15, 56–67 (2011).

46. Rademacher, L. et al. Dissociation of neural networks for anticipation and consumption of monetary and social rewards. NeuroImage 49, 3276–3285 (2010).

47. Chevallier, C., Kohls, G., Troiani, V., Brodkin, E. S. & Schultz, R. T. The social motivation theory of autism. Trends Cogn. Sci. 16, 231–239 (2012).

48. Haggarty, C. J., Malinowski, P., McGlone, F. P. & Walker, S. C. Autistic traits modulate cortical responses to affective but not discriminative touch. Eur. J. Neurosci. n/a, (2019).

49. Walker, S. C. & McGlone, F. P. The social brain: neurobiological basis of affiliative behaviours and psychological well-being. Neuropeptides 47, 379–393 (2013).

50. Fischer, A. G. & Ullsperger, M. An Update on the Role of Serotonin and its Interplay with Dopamine for Reward. Front. Hum. Neurosci. 11, (2017).

51. World Medical Association. World Medical Association Declaration of Helsinki: ethical principles for medical research involving human subjects. JAMA 310, 2191–2194 (2013).

52. Fridlund, A. J. & Cacioppo, J. T. Guidelines for human electromyographic research. Psychophysiology 23, 567–89 (1986).

53. Bates, D., Maechler, M., Bolker, B. & Walker, S. lme4: Linear mixed-effects models using Eigen and S4. (2014).

54. R Core Team. R: A language and environment for statistical computing. (R Foundation for Statistical Computing, 2019).

55. Judd, C. M., Westfall, J. & Kenny, D. A. Treating stimuli as a random factor in social psychology: A new and comprehensive solution to a pervasive but largely ignored problem. J. Pers. Soc. Psychol. 103, 54–69 (2012).

56. Delorme, A. & Makeig, S. EEGLAB: an open source toolbox for analysis of single-trial EEG dynamics including independent component analysis. J. Neurosci. Methods 134, 9–21 (2004).

